# Stage-specific phenotypic and transcriptional alterations in keratinocytes exposed to acute and chronic blue light

**DOI:** 10.1101/2024.11.21.624322

**Authors:** Paulo Newton Tonolli, Suely Kazue Nagahashi Marie, Sueli Mieko Oba-Shinjo, Leonardo Vinicius Monteiro de Assis, Maurício S. Baptista

**Affiliations:** Department of Biochemistry, Institute of Chemistry, University of Sao Paulo, Sao Paulo, Brazil; Department of Neurology, Laboratory of Molecular and Cellular Biology, LIM15, Faculdade de Medicina FMUSP, Universidade de Sao Paulo, Brazil; Institute of Neurobiology, Center of Brain Behavior & Metabolism, University of Lübeck, Germany; University Hospital Schleswig-Holstein, Campus Lübeck, Lübeck, Germany

**Author notes:** Corresponding author: Maurício S. Baptista. shared authorship (Leonardo Vinicius Monteiro de Assis, Maurício S. Baptista).

## Abstract

Despite evidence that visible light (VL) has similar effects on human skin as those of UVA, VL is often viewed as harmless. High SPF sunscreen prevents erythema but can lead to overexposure to UVA and VL, with unknown consequences. To explore the impact of chronic blue light exposure, we irradiated (50 J/cm², λ= 408 nm, three times a week) human immortalized keratinocytes under acute (3 irradiations), intermediate (14 irradiations), and chronic (42 irradiations) blue-light exposure, monitoring phenotypic and gene expression changes. Chronically exposed keratinocytes exhibit increased nuclei area, chromatin alterations, higher proliferation, and apoptosis resistance, mirroring the consequences of chronic UVA exposure. While acute exposure upregulated keratinization and downregulated tissue repair and apoptosis genes, chronically exposed cells had upregulated genes involved with energy metabolism and oxidative phosphorylation and downregulated genes were enriched for immune and inflammatory responses. Specific transcriptional factors were identified in the acute and chronic stages, some of them had been associated with UVB exposure. We identified some changes in chronically irradiated keratinocytes similar to the malignant transformation, emphasizing the need for further research on the long-term impacts of blue light exposure on human skin.

## INTRODUCTION

Solar radiation reaching the Earth surface consists of ultraviolet (UV) radiation (280–400 nm, ca. 2% of the solar spectra), visible light (VL, 400–750 nm, ca. 47%), and infrared radiation (750–2500 nm, ca. 51%) (1). In recent decades, research has mainly concentrated on UV wavelengths, particularly UVB (280–320 nm) and UVA (320–400 nm), because of their role in causing DNA damage and heightening oxidative stress. These factors are well-known to elevate the risk of skin cancer (1–4). However, UV radiation also has beneficial effects. For instance, vitamin D synthesis depends on cutaneous exposure to UVB radiation (5, 6). Furthermore, exposure to solar radiation has been linked to reduced blood pressure (7). Experimental studies have also demonstrated that low doses of UVB can reduce weight gain and liver steatosis in mice on a high-fat diet (8, 9) and affect the hypothalamus-pituitary-adrenal axis (10).

Unlike UV radiation, visible light is generally considered safe for the skin. Most sunscreens are formulated to protect against UVB and UVA radiation but offer little to no protection against VL and infrared radiation. Ironically, while sunscreen prevents erythema by blocking UV radiation, it can increase overall solar exposure, as it gives a false sense of protection. This may lead to prolonged exposure to VL and potentially skin damage (1). Although VL causes less direct DNA damage than UV radiation, it increases oxidative stress. This is mainly because the endogenous photosensitizers, which are responsible for the effects of UVA radiation in human skin, also absorb VL (11). Indeed, the effects of UVA and VL can be mitigated by incorporating antioxidants such as vitamins C and E into skincare routines (12, 13). The VL spectrum includes red (625–740 nm), orange (590–625 nm), yellow (565–590 nm), green (500–565 nm), blue (450–485 nm), and violet (400–450 nm). Among these, blue light has been extensively studied for its biological effects, including its role in human cell proliferation and differentiation, increased oxidative stress in keratinocytes (14), the systemic release of β-endorphin (15), hair follicle growth in ex vivo models (16), as well as anti-inflammatory (17), and vasorelaxation effects (18). The oxidative stress induced by VL is highly dependent on its wavelength, with violet and blue light being the primary contributors to reduced cellular viability, DNA damage, and lysosomal dysfunction (19). Pre-exposure of human keratinocytes to UVA radiation further sensitizes cells to VL, as it triggers lipofuscin formation. Lipofuscin acts as a visible light photosensitizer, amplifying oxidative stress and increasing DNA damage (20, 21). These findings suggest that visible light-induced skin damage is mediated by the photoexcitation of endogenous photosensitizers, such as lipofuscin (19, 21, 22). Moreover, blue light exposure is also known to inhibit DNA repair (23). Unlike the substantial research on the long-term effects of UVA exposure (24), there is a lack of studies examining the chronic impacts of blue light exposure. Fortunately, products that provide adequate blue light protection are already available (25, 26).

To expand the knowledge into the deleterious effects of excessive blue light exposure, we investigated the impacts of acute and chronic blue light stimulation on immortalized human keratinocytes. Phenotypic changes included increased nuclear size, alterations in chromatin structure, enhanced cell proliferation, and resistance to UVA-induced apoptosis after chronic blue light irradiation. Transcriptome analysis revealed significant early and late stage-specific transcriptional alterations. We identified transcription factors associated with acute and chronic blue light exposure. Interestingly, several of these transcription factors are also affected by UV radiation, which suggests a shared mechanism between UV and blue light. Our findings offer a comprehensive molecular framework to understand the long-term effects of blue light on skin cells.

## MATERIAL AND METHODS

### Cell culture

Human skin immortalized keratinocyte cells (HaCaT) (27) were cultivated in Dulbecco modified Eagle medium (DMEM), supplemented with 10% fetal bovine serum (FBS), 4 mM L-glutamine, 100 U/mL penicillin and 100 pg/mL of streptomycin, in 5% CO2 at 37°C.

### Light source and irradiation protocols

HaCaT cells (5 x 10⁵ cells per 100 mm plate) were exposed to a blue light LED source (λ = 408 nm, irradiance = 0.28 mW.cm^-2^) developed by the Optics and Photonics Research Center at the Institute of Physics, University of São Paulo. The cells were irradiated in phosphate-buffered saline at an irradiation of 50 J/cm² three times a week for up to 14 weeks. Light dose was constant through the experiment and chosen to provide 50% cell death in HaCaT cells. The dose was calculated to be physiologically relevant, based on comparing the emission intensity of blue light LED source (λ = 408 nm) profile with the standardized solar spectrum Air Mass 1.5 spectrum (AM 1.5), a reference spectrum of sun irradiance at sea level for typical latitudes of most major cities with the sun at about 48° from zenith (irradiance = 0.13 mW.cm^-2^). After each irradiation, the culture medium was reintroduced until the next session. Cells were passaged every 3-5 days. To assess resistance to UVA radiation, 1.5 x 10³ cells per well were plated in a 48-well plate. After 12 hours, the cells were irradiated with different irradiation doses of UVA (6 and 12 J.cm^-2^) using a LED source (λ = 365 nm, BioLambda) at 30°C.

### Quantification of Nuclear Area

Approximately 2 x 10⁵ cells were seeded on coverslips in 60 mm. After 24 hours, the cells were rinsed with PBS and fixed with 4% paraformaldehyde (w/v) for 10 minutes. The cells were then washed three times for 5 minutes each with PBS. Cells were permeabilized by incubating with 0.1% Triton X-100 solution for 10 minutes at 37°C. Following another PBS wash, the cells were stained with DAPI (4’,6-diamidino-2-phenylindole) by mounting the coverslips with Prolong-Gold Antifade containing DAPI (ThermoFisher Scientific, P36935). The slides were then analyzed using a Zeiss LSM 510 Meta confocal microscope, with excitation at 351 nm to visualize the DAPI-stained nuclei. Micrographs were converted to 8-bit images, and the threshold was adjusted using polygonal selection to delineate the nuclear area. Nuclear area measurements were performed on 100 nuclei from fluorescence micrographs stained with DAPI.

### BrdU Cell Proliferation Assay

Approximately 5 - 10 x 10^3^ cells were plated in a 96-well plate. After 12 hours, the cells were incubated with BrdU solution in 10% DMEM for four hours in a 5% CO₂ atmosphere at 37°C. BrdU detection was performed using an ELISA assay (absorbance read at 490 nm) following the manufacturer’s protocol, using the BrDU Cell Proliferation Assay Kit (Cell Signaling Technology).

### Determination of Apoptosis

Apoptosis was assessed 18 hours after UVA irradiation (λ = 365 nm, 6 and 12 J.cm^-2^) using Annexin-V/propidium iodide staining (ThermoFisher Scientific) and analyzed by flow cytometry according to the manufacturer’s instructions.

### RNA sequencing

RNASeq Libraries were prepared using a QuantSeq 3’ mRNA-Seq Library Prep Kit-FWD (Lexogen, Vienna, Austria) with 1μg RNA. The library concentration was measured by Qubit Fluorometer and Qubit dsDNA HS Assay Kit (Applied Biosystems), and the size distribution was determined using an Agilent D1000 ScreenTape System (Agilent Technologies). Sequencing was performed on NextSeq 500 platform at the NGS facility core SELA. We used STAR to align The sequencing data was aligned to the GRCh38 version of the human genome using STAR, and the bamsort tool, from biobambam2, for downstream processing of the BAM file, including merging, sorting and marking of duplicates. To count the number of reads that overlap each gene, we used featureCounts.

### Bioinformatic analysis

DESeq2 (28) was used to analyze differentially expressed genes (DEGs) from raw count data. To identify genes influenced by time or blue-light exposure, a likelihood ratio test (LRT) was conducted under standard conditions (padj < 0.1), considering the influence of time and blue-light stimulation. Unsupervised clustering was then performed using the "complete" method with Euclidean distance to categorize the DEGs. Pairwise comparisons were also carried out using the Wald test with standard parameters (padj < 0.1). Subsequent enrichment analyses were conducted using gProfiler.

### Statistical analyses

Analyzes were performed in either RStudio (v. 4.2.1) or GraphPad Prism (v. 10). Pair-wise comparisons were made using Student’s t test with Welch Correction. When comparing two factors, Two-Way ANOVA followed by Tukey post-test was used. A p-value < 0.05 was considered significant.

### Data availability

The processed data derived from the RNA sequencing is presented in Table S1. Additional data that support the findings in this manuscript can be obtained from the corresponding author upon reasonable request.

## RESULTS

### Chronic blue light irradiation alters keratinocyte morphology, proliferation, and apoptosis resistance

We investigated the effects of acute and chronic blue light irradiation (λ = 408 nm, 50 J/cm², three times per week) on immortalized human keratinocyte (HaCaT) cells, by using a protocol involving repeated blue light exposure. Normal HaCaT (control) cells were chronically exposed to blue light (50 J/cm^2^, three times a week for up to 14 weeks). We kept control and irradiated cells with the same regime of passage and evaluated both the controls and the irradiated cells with the same number of passages. We classified the cells as acute (3 irradiations), intermediate (14 irradiations) and chronically (42 irradiations) exposed to blue light. Morphological differences between chronically blue-light-exposed cells and control cells were observed starting from the 19^th^ irradiation. Control cells displayed an epithelial-like morphology with uniform size and shape (Figure 1A, panels a, c). Note that high passage cells have not marked cell phenotype alterations. In contrast, blue-light-exposed cells showed significant morphological changes, including the presence of giant cells with multiple and irregularly shaped nuclei and altered nucleoli (Figure 1A, panels b, d). Blue-light-exposed cells exhibited almost double the nuclear area compared to control cells (p < 0.0001; Figure 1B). Under light microscopy, normal nuclei exhibit fine and uniformly distributed chromatin, whereas malignant cells display coarse or clumped chromatin with an irregular distribution, forming transparent regions in the nucleus center. Such features were identified in blue light-exposed cells compared to the control (Figure 1C). Furthermore, blue-light exposure increased cell proliferation, as indicated by BrdU incorporation (Figure 1D). Blue-light-exposed cells also resisted to UVA-induced apoptosis (λ = 365 nm, 12 J/cm²), showing a significant reduction in the total percentage of apoptotic cells compared to control cells (Figure 1E). These phenotypic changes are very similar to those observed in HaCaT cells that experienced a malignant transformation due to a chronic UVA exposure protocol (24). Taken altogether, our findings show that chronic blue light induces several changes in keratinocytes, indicating a step toward malignization.

**Figure 1:**
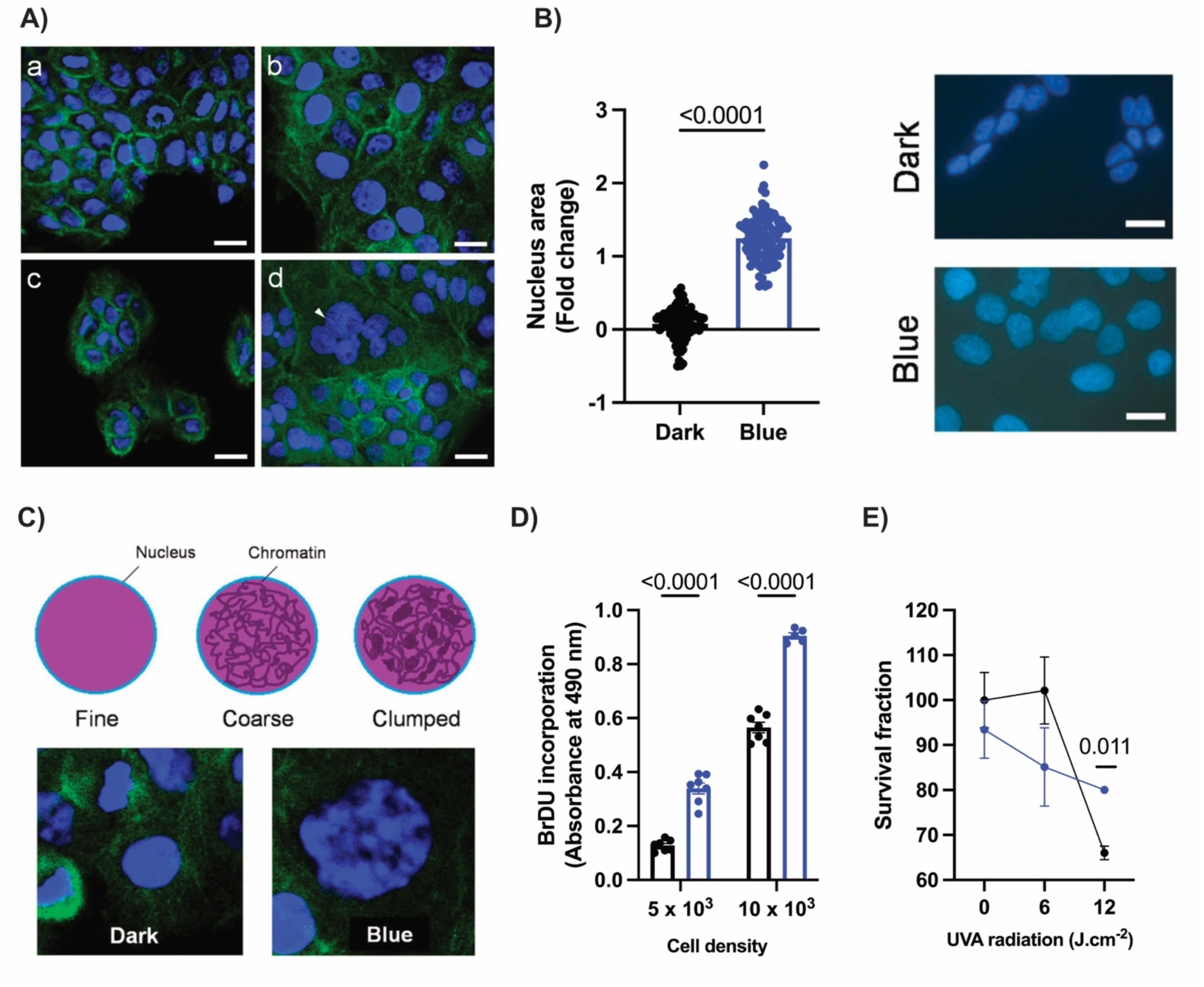
Phenotypic evaluation of chronic blue-light exposed cells. A) Representative figures of control (a and c) and chronic blue-light exposed (b – d) cells. Scale bar = 20 μm. (a) Control (dark) cells, with a typical compacted organization. (b) Chronic blue light treated cells, showing giant cells and increased nuclei size. (c) Control (dark) cells with mononucleated cells. (d) Chronic blue light treated cells depicting multinucleated cells (white arrows). Scale bar = 20 μm. B) Quantification of nucleus area. An unpaired Student’s t-test was used. Representative images are depicted. Scale bar = 20 μm. C) Representative chromatin visualization of control and blue-light exposed cells. D) Proliferative capacity of control and chronic blue-light exposed cells measured by BrdU incorporation. E) Percentage of surviving chronic blue-light exposed cells and controls with increasing UVA dose (representative of two independent experiments).

### Stage-dependent transcriptome modulation in human keratinocytes following acute, intermediate, and chronic blue light irradiation

To evaluate the temporal dynamics involving blue-light exposure in keratinocytes, we assessed the transcriptome of cells exposed to three different conditions: acute, intermediate, and chronic exposure, corresponding to 3, 14, and 42 exposure to blue light, respectively. Principal component analysis (PCA) revealed distinct separations between the groups exposed to varying stages of blue light. Notably, a clear distinction was observed between the blue light-exposed and control groups in both the acute and chronic exposure conditions, while the separation was less pronounced in the intermediate group (Figure 2A).

**Figure 2:**
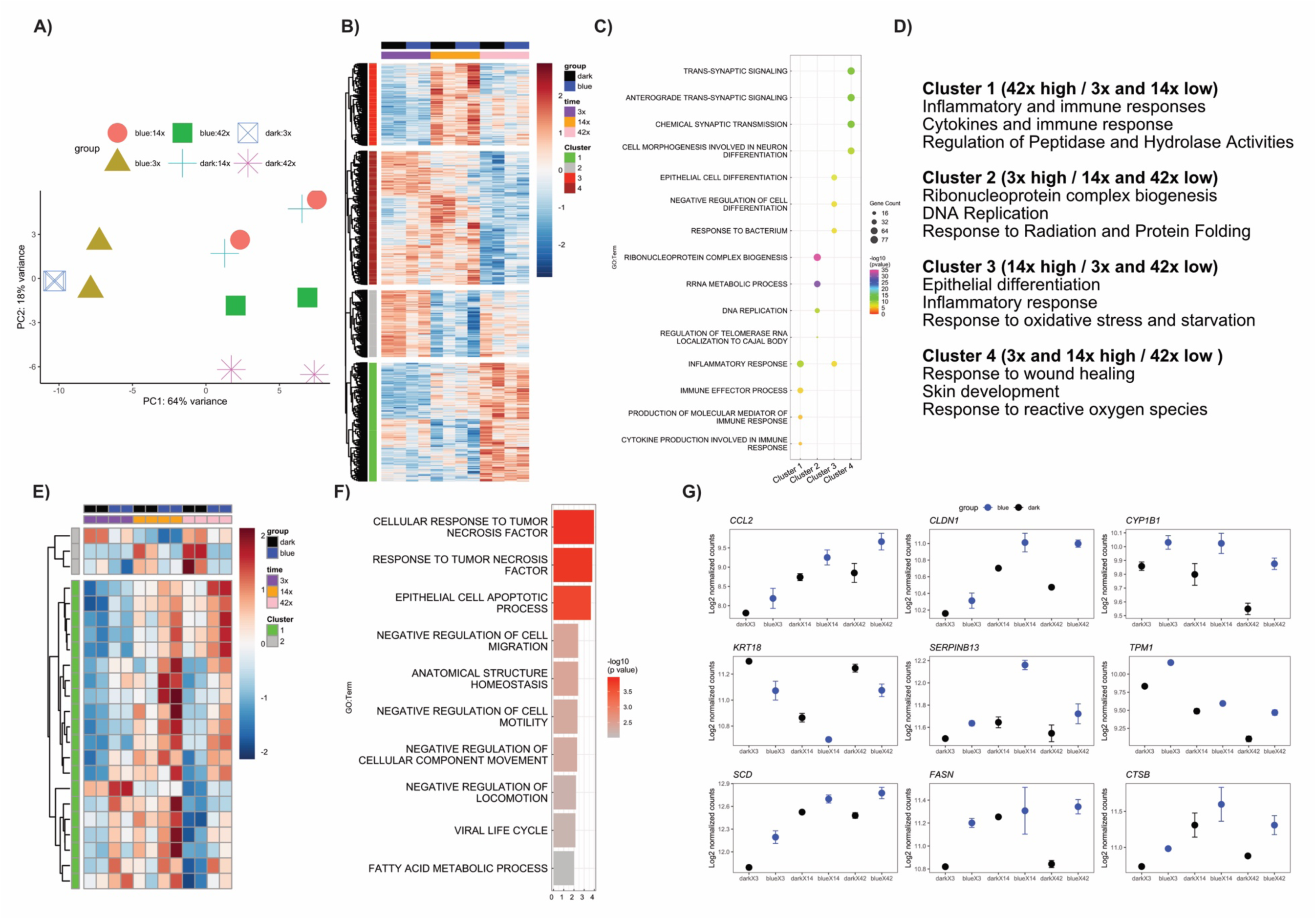
Identification of differentially expressed genes (DEGs) in response to time (passage effect) and blue light stimulation. A) Principal component analysis (PCA) of all samples. B) The heatmap depicts the DEGs that show a time effect, identified by DESEq2 using the likelihood ratio test (LRT, padj < 0.1, fold change ± 0.2). C) Enrichment analysis was performed in each cluster using gProfiler. D) Summary of the biological processes identified in C. E) The heatmap depicts the DEGs that show a blue light effect, identified by DESEq2 using the likelihood ratio test (LRT, LRT, padj < 0.1). F) Enrichment analysis was performed on the DEGs identified in E. G) Representative genes identified in E and F are depicted.

An important aspect of our experimental design is the temporal nature of blue light irradiation, with cells exposed to different light conditions over several weeks. Consequently, cells across the experiment accumulated an increasing passage number, which resembles an aging process. To analyze these effects, we employed DESeq2 using the Likelihood Ratio Test (LRT), incorporating both blue light exposure and time as factors in the full model. This approach allowed us to capture the effects of time on gene expression. The reduced model focused on the effects of time (cellular passage) alone to explain variance in the transcriptome data. Our analysis revealed a significant effect of the number of passages across the experiment, identifying 2,815 differentially expressed genes (DEGs) (padj < 0.1 and fold change ± 0.2, Table S1). Unsupervised clustering of these DEGs revealed four distinct gene expression patterns (Figure 2B). Enrichment analyses of these clusters highlighted unique biological processes associated with different stages of the experimental protocol. It is important to note that no changes in phenotype were detected in any of the control groups. Therefore, the changes detailed below occur under conditions of cellular homeostasis.

Processes associated with the time effect (number of passage) were enriched and clustered. Genes associated with ribonucleoprotein complex biogenesis, DNA replication, telomere organization, response to radiation, and protein folding were upregulated in the acute exposed group (3 irradiations) but significantly downregulated in subsequent stages (Cluster 2). In the intermediate stage (14 irradiations), significant upregulation was observed in genes associated with epithelial differentiation, inflammatory response, oxidative stress, and starvation (Cluster 3). In contrast, genes involved in wound healing, skin development, and response to reactive oxygen species were upregulated in both the acute and intermediate stages but showed marked downregulation in the chronic stage (Cluster 4). Importantly, the chronically exposed group (42 irradiations) had upregulated genes enriched, which were related to inflammatory and immune responses, cytokine activity, and regulation of peptidase and hydrolase activities (Cluster 1, Figure 2B – D, Table S1). The gene expression profile of the high passage group likely renders these cells more capable of enduring photoinduced damage compared with the control and cells submitted to a smaller amount of passage.

In the DESEq2 reduced model, we also evaluated genes specifically responding to blue light irradiation and identified a subset of 23 differentially expressed genes (DEGs) that exhibited consistent responses throughout the experimental protocol. These DEGs were enriched for processes related to tumor necrosis factor signaling, cellular apoptosis, cellular motility, and fatty acid metabolism. Notably, the small number of DEGs identified across the different irradiation series suggests that the effects of blue light irradiation are stage-dependent (Figure 2E-G, Table S1).

### Stage-dependent transcriptome changes in blue light-exposed human keratinocytes

To further explore this stage-dependent regulation, we performed pairwise analyses for each stage using DESEq2, which enabled us to pinpoint stage-specific transcriptome alterations. Acute blue light irradiation increased and decreased the expression of 38 and 15 genes, respectively (padj < 0.1, Table S1). Upregulated genes were enriched in processes such as intermediate filament organization, keratinization, cellular differentiation, and interleukin-1 biosynthesis. Downregulated genes were associated with protein kinase activity, positive regulation of apoptosis, interleukin-17 biosynthesis, wound healing, and metabolism of nitrogen- reactive species (Figure 3A–B, Table S1).

**Figure 3:**
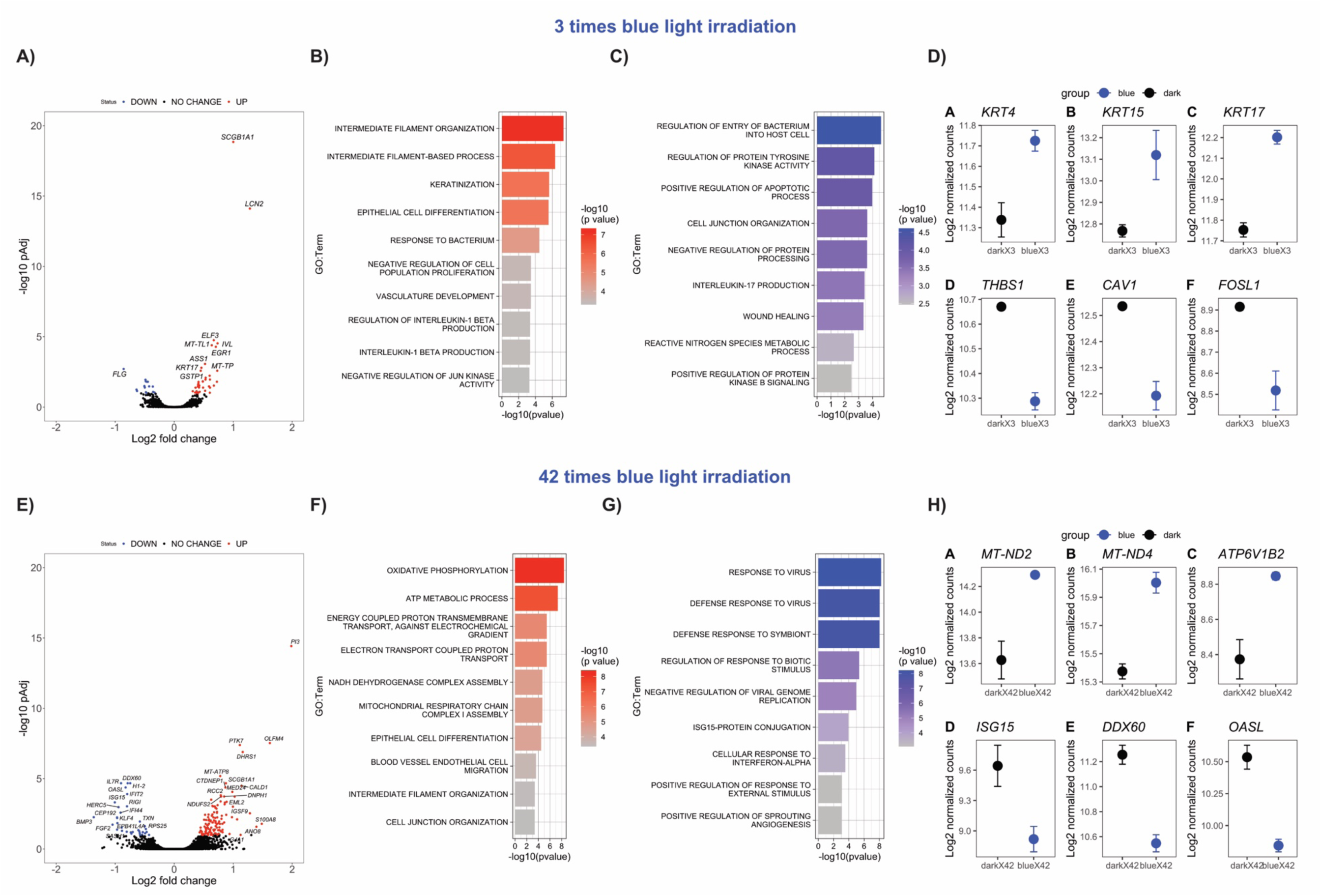
Identification of differentially expressed genes (DEGs) in response to 3 or 42 times blue light stimulation. A and E) Volcano plots depict the up- and downregulated DEGs identified in DESEq2. B, C, F, and G) Enrichment analyses were performed on the identified DEGs using gProfiler. D and H) Representative genes identified in the previous analyses are shown.

In the intermediate stage (14 irradiations), we observed a lack of response to blue light compared to the control. This unexpected finding prompted us to re-investigate the gene signatures influenced by time (i.e., cellular passage), rather than the blue light, identified in Cluster 3 (Figure 2B). In this stage, several genes associated with oxidative stress (*SESN3*, *ULK1*, *PRKAA2*, *BMPR2*, and *DAP*) and starvation (*TXNIP*, *NFE2L1*, *GPX2*, *PRDX5*, and *SESN3*) were identified. Considering the lack of response to blue light in the intermediate group, we suggest that the resistance phenotype observed in keratinocytes is likely due to their enhanced ability to manage oxidative stress and maintain cellular homeostasis through processes like autophagy and antioxidant defense, which are critical for the blue-light induced effects (19–21).

Interestingly, in the chronic stage, the resistance phenotype to blue light was lost, resulting in increased detection of DEGs. This is in accordance with the gene expression profiling identified in the temporal effects (passage effects, see above). A total of 115 upregulated and 34 downregulated genes were identified (padj < 0.1, Table S1). Upregulated genes were enriched in energy metabolism processes such as oxidative phosphorylation, ATP metabolism, mitochondrial respiration, and cell differentiation. In contrast, downregulated genes were associated with innate immune response, response to interferon-alpha, and response to external stimuli (Figure 3E – H, Table S1).

### Identification of transcriptional factors associated with acute and chronic blue-light exposure

Considering the transcriptome alterations evoked by blue light in both acute and chronic treatments, we used the ChEA3 prediction tool (29) to identify potential transcriptional factors associated with the stage-dependent gene signature (Figure 4A). A clear stage-specific TF signature was identified as being consistent with the stage-specific transcriptome signature. During acute blue light irradiation (3 irradiations), several transcriptional factors were significantly upregulated, except for FOSL1 (Figure 4 B-C). The molecular function of these factors is described next. IRF1 is a key regulator of immune responses and is involved in apoptosis and stress response (30). EGR1 is an immediate-early gene that responds rapidly to stress and growth factors, being an important regulator of cell growth, differentiation, and apoptosis (31, 32). ELF3 is crucial for regulating epithelial cell differentiation (33). FOSL1, part of the AP-1 transcription factor complex, is known to regulate cell proliferation, differentiation, and survival, being rapidly induced by stress and inflammatory signals (34). Lastly, ZNF704 is a member of the zinc finger protein family involved in differentiation, metabolism and apoptosis, and it has been suggested to be an oncogene (35, 36). SOX4 is a critical regulator of cell survival and apoptosis that is also considered an oncogene (37). Importantly, most identified TFs are associated with stress response against damaging factors, including oxidative stress.

**Figure 4:**
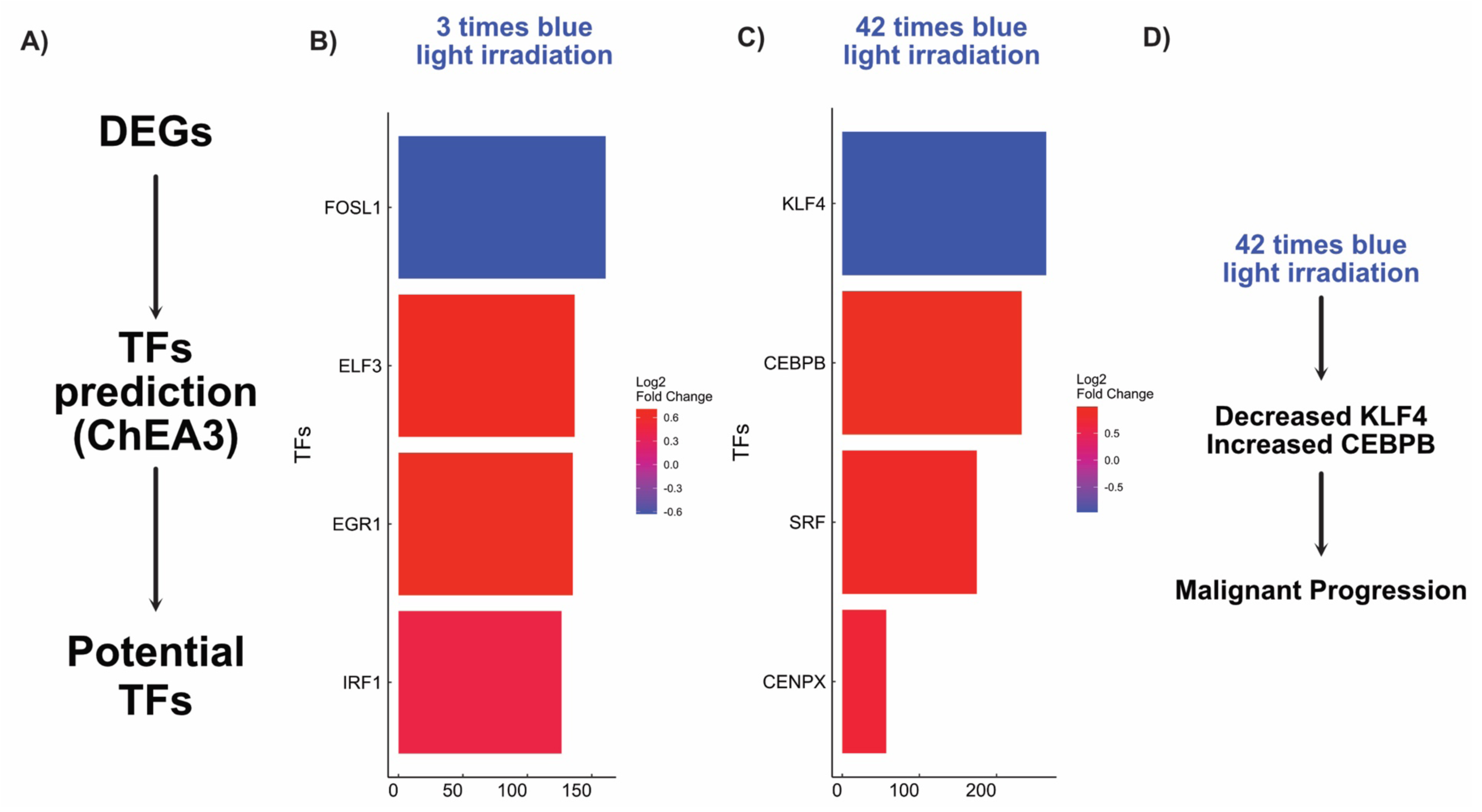
Identified putative transcriptional factors (TFs) in response to acute or chronic blue light stimulation. A) The data analysis pipeline description used to identify TFs using ChEA3 is shown. B and C) Top 4 identified TFs as being up- (red) or down-regulated (blue) in the acute or chronic blue light stimulation. D) Overview of the conclusions drawn for the chronic blue-light stimulated group.

In contrast, chronic blue light irradiation (42 irradiations) altered different transcriptional factors. CENPX is involved in DNA repair and chromosomal stability, indicating a long-term adaptation to sustained DNA damage (38). SRF regulates cytoskeletal dynamics, cell growth, and survival (39). CEBPB plays significant roles in inflammatory responses, cell survival, and differentiation, potentially indicating a shift towards a sustained inflammatory state and repair mechanisms (40, 41). Finally, KLF4 is involved in maintaining cellular homeostasis, stress responses, differentiation, and apoptosis, promoting cellular survival and adaptation over time (42–44). Importantly, our findings show that blue light effects on keratinocytes are stage-dependent and are driven by a different set of transcriptional factors.

## DISCUSSION

This study investigated the effects of acute and chronic blue light irradiation on human keratinocytes, revealing significant stage-dependent alterations in cellular morphology and transcriptome alterations. Chronic exposure increased nuclear size, chromatin alterations, enhanced proliferation, and resistance to UVA-induced apoptosis. Transcriptome analyses across different exposure stages revealed distinct gene expression patterns, with early stages marked by upregulation of genes related to keratinization and downregulation of those involved in immune and inflammatory responses. Notably, cells exposed to intermediate levels of blue light exhibited a resistance phenotype, which was lost in chronically blue-light-exposed cells. In these cells, oxidative phosphorylation genes were upregulated, while immune response genes were downregulated, indicating a potential shift towards energy generation.

The phenotypic changes observed, such as increased nuclear size and chromatin reorganization, are consistent with features of malignant transformation, as previously reported in HaCaT cells (24, 45). These nuclear alterations, including irregular shapes and chromatin clumping, suggest shifts in gene expression through chromatin remodeling and histone modifications, which are key processes in DNA replication and repair (46, 47). Increased proliferation and resistance to apoptosis further support this malignization, aligning with established cancer hallmarks (48), and have also been observed in chronically UVA-exposed cells (24).

While our study focused on blue light, we also accounted for the influence of cellular aging in culture, a critical factor in long-term experiments. By tracking changes over time, we identified 2,815 DEGs and used unsupervised clustering to pinpoint stage-specific transcriptomic alterations linked to aging. It is unlikely that these analyses are influenced by blue light since they include the group exposed to blue light. Importantly, these findings show a highly dynamic transcriptome regulation dependent on the time the cells are in culture, which is a critical factor that should be considered in long-term cell culture studies. On the other hand, one limitation is the use of immortalized HaCaT cells, which bypass natural senescence and apoptosis, making them a model more suitable for studying certain chronic effects, though not entirely representative of physiological cell aging. A general observation regarding the cell alterations with an increasing number of passages is that no noticeable phenotypic changes occurred. Yet, the gene expression profile changes substantially, highlighting a reduction in the expression of key regulatory genes essential for protecting cells against oxidative stress. In summary, aged cells appear to be more resilient to conditions that induce oxidative imbalance, such as exposure to blue light, which can represent an important step toward malignant transformation.

When focusing on the effects of blue light exposure, we found that changes were distinctly stage-specific. Our bioinformatic analyses revealed that acute blue light exposure led to upregulation of genes involved in keratinization, filament organization, and interleukin-1 signaling, while genes associated with apoptosis and wound healing were downregulated. However, no blue light effect was observed after 14 exposures (intermediate stage). Interestingly, in this stage, the cells had a particular gene signature, independently of the blue-light stimulation, but rather associated with the cellular moment related with oxidative stress, which may have contributed to the observed effects. Importantly, such resistance phenotype was lost in chronically exposed cells. In this stage, cells exhibited increased expression of oxidative phosphorylation genes and decreased expression of innate and inflammatory response genes. These findings strongly suggest an altered energy metabolism profile in chronic blue light-exposed cells.

A study conducted on *Drosophila* maintained under a 12-hour light and 12-hour dark cycle, in which blue light was administered at significantly lower irradiance levels compared to our experiment, found that older flies exhibited a greater susceptibility to blue light due to diminished mitochondrial capacity (49). Despite the challenges inherent in comparing our experiment with the earlier study (49), it is noteworthy that under elevated levels of blue light irradiation (our findings), an adaptive mechanism is triggered that enhances resistance to the effects of blue light. A recent study utilizing reconstructed skin subjected the explants to low levels of blue light irradiation, similar to the doses encountered in electronic devices. Although the irradiance remains significantly low in comparison to our conditions, this study demonstrated that, even under such reduced irradiance, blue light-induced an inflammatory response (e.g., elevated IL6, STAT3, and PPAR expression) as well as a pigmentary response (50).

Skin cells express a diverse array of light-sensitive molecules known as opsins (1, 51). Numerous studies have identified that the effects of blue and violet light on both skin and non-skin cells are dependent on the presence of panopsin (OPN3) (16, 52–56). Importantly, most of these investigations concentrated on scenarios involving acute stimulation, whereas the present study examines chronic stimulation. Notably, under the experimental conditions established in this study, OPN3 levels remained unchanged in response to either acute or blue light stimulation (Table S1).

Transcriptional changes in the acutely blue light-exposed cells were associated with specific transcription factors, including *FOSL1*, *IRF1*, *EGR1*, *ELF3*, *ZNF704*, and *SOX4*. The *FOSL1* was the only one downregulated. A shared regulatory mechanism was identified between blue light and UV radiation for some of these factors. For example, reduced *FOSL1* expression was reported in radiation-induced neoplastic transformation of human *CGL1* cells (57). Additionally, UV radiation increases IRF3 and NF-κB activity in melanocytes, contributing to PD-L1 expression and immune suppression (58). Similarly, UV exposure increases *STAT1* and *IRF1* levels in keratinocytes, promoting pro-inflammatory cytokine production (59). *EGR1*, another transcription factor linked to UV exposure, was particularly elevated in p53 knockout cells, suggesting that p53 regulates its expression (60). Moreover, UVC, UVB, and UVA radiation exposure have been shown to rapidly induce EGR1 expression (61). As for ELF3 and ZNF704, their roles in response to UV radiation remain unexplored, but they emerge as novel markers of blue light exposure.

Chronic blue light exposure significantly alters the expression of four key transcription factors in keratinocytes. Our findings provide novel evidence linking chronic blue light stimulation with the dysregulation of CENPX and SRF. Consistent with previous studies using UV radiation, our findings show that chronic blue light exposure alters the expression pattern of CEBPB and KLF4. Collectively, we suggest these changes contribute to a substantial alteration in the keratinocyte transcriptome, potentially promoting conditions that result in malignancy. Our data corroborates the literature. CEBPB is critical for keratinocyte survival in response to carcinogenic treatment (62, 63). Specifically, CEBPB is essential for Ras-induced skin tumorigenesis, as evidenced by the resistance of CEBPB knockout mice to skin tumors initiated by carcinogen treatment (62). CEBPB plays a crucial role in supporting the survival of Ras-mutant tumor precursor cells by regulating p53 levels and preventing apoptosis in keratinocytes (63). Furthermore, CEBPB has been shown to repress p53 to enhance cell survival after DNA damage, and the loss of CEBPB increases the effectiveness of chemotherapeutic agents (64). A study has highlighted the significant role of CEBPB in regulating the p53-mediated apoptotic response in keratinocytes, thereby increasing cell survival and susceptibility to UVB-induced skin cancer. Interestingly, mice lacking CEBPB in the epidermis were resistant to UVB-induced skin cancer (65). However, it is noteworthy to highlight that repeated UVB radiation (5 irradiations, 12 J/cm²) followed by a week of recovery reduced CEBPB expression (66). In contrast, KLF4 knockdown increases cellular growth and migration in human keratinocytes (67). Chronic UVB radiation also results in a progressive decrease in KLF4 expression, which is downregulated in human skin tumors (67). Extensive evidence supports the role of KLF4 as a tumor suppressor gene (68). Importantly, our study provides strong evidence of KLF4 downregulation in response to blue light, consistent with previous studies focused on UV radiation.

Several lines of evidence from both morphological and transcriptional analyses suggest a progression toward malignancy in chronically blue light-exposed cells. Importantly, these cells exhibit some signs of malignant transformation, indicating that additional factors beyond blue light exposure are required for a complete transformation. This aligns with current cancer development models, which suggest that mutations in multiple genes, often in a stepwise process, are necessary for complete malignant transformation (69–71). Notably, HaCaT cells, which carry mutated forms of p53 (72), imply that further genetic or cellular alterations are needed to drive malignancy. Nevertheless, our findings highlight the potential of blue light to induce substantial cellular changes that, when combined with mutations in key tumor suppressor genes and oncogenes, may serve as a critical driver of malignant transformation. Additional experimental models, especially those involving human keratinocytes with mutated tumor suppressor genes, are essential to investigate this further. Crucially, our results emphasize the potentially harmful effects of blue light on human skin, emphasizing the need for continued research.

## CONCLUSION

In summary, our study provides a comprehensive view of the stage-specific effects of blue light on human keratinocytes, both at the phenotypic and transcriptomic levels. These findings shed light on the mechanisms underlying blue light-induced cellular damage and suggest shared pathways with UV radiation exposure. Notably, the level of blue light exposure in our study is much higher than that emitted by digital devices, emphasizing that our findings are more relevant to blue light from sunlight. This highlights the need for further research to fully understand the long-term consequences of blue light from sun exposure on skin health. From a public health perspective, our results contribute to the increasing body of evidence showing the damaging and deleterious effects of blue light on the skin. Protective measures, such as sunscreens that block visible light, should be further investigated in human studies (25).

## ARTIFICIAL INTELLIGENCE STATEMENT

While preparing this study, the authors utilized ChatGTP to enhance readability and language. Following this, they thoroughly reviewed and edited the content as necessary, taking full responsibility for the publication’s content.

## ACKNOWLEDGMENTS AND FUNDING

The author(s) declare that financial support was received for the research, authorship, and/or publication of this article. Baptista, M received grants From Sao Paulo Research Foundation (2022/13066-9, 2021/08521-6, and 2013/07937-8). de Assis, LVM received a startup grant and funding from the Local Control of Thyroid Hormone Action (LOCOTACT) Consortium (project ID 424957847-TRR 296) and from a research grant for basic science from the European Thyroid Association (ETA 2023). The authors thank Jan Schlothauer and Felipe Ravagnani for their technical assistance and support.

## AUTHOR’S CONTRIBUTION

Paulo Newton Tonolli: conceptualization, methodology, data acquisition, data curation, manuscript writing and review. Suely Kazue Nagahashi Marie: data curation; manuscript review. Sueli Mieko Oba-Shinjo: data curation; manuscript review; Leonardo Vinicius Monteiro de Assis: supervision, conceptualization, manuscript writing and review. Maurício S. Baptista: Funding, supervision, conceptualization; manuscript writing and review.

